# Comparative Analysis of Two-Dimensional and Three-Dimensional MRI Radiomics Models for Predicting Molecular Biomakers in Breast Cancer

**DOI:** 10.1101/2025.11.26.690866

**Authors:** Xinyu Wang, Xiaodong Liu, Zhiqing Wang

**Author notes:** **Corresponding Author:** Xinyu Wang, **Corresponding Email:**.

## Abstract

**Background:** Breast cancer (BC) incidence continues to rise globally, with diverse clinical presentations that remain challenging to predict.

**Purpose:** To compare the predictive performance of conventional 2D DCE-MRI-based radiomics models with advanced 3D DCE-MRI-based radiomics models for determining estrogen receptor (ER), progesterone receptor (PR), human epidermal growth factor receptor 2 (HER-2), and Ki-67 expression in breast cancer patients.

**Material and Methods:** This study included 385 patients with histologically confirmed breast cancer. Semi-automatic segmentation was applied to delineate regions of interest (ROIs) on DCE-MRI images, and radiomic features were extracted using Intelligence Foundry (IF) 3.1. Feature selection involved single-value filtering, correlation analysis, outlier optimization, normalization, and least absolute shrinkage and selection operator (LASSO) regression. Logistic regression was used for statistical analysis, and model performance was evaluated using area under the receiver operating characteristic curve (AUC), decision curve analysis (DCA), and DeLong’s test for comparing 2D and 3D models.

**Results:** The 3D models consistently outperformed 2D models across all biomarkers. In the validation set, the 3D models achieved AUCs of 0.884 for ER, 0.861 for PR, 0.898 for HER-2, and 0.886 for Ki-67, while the 2D models yielded AUCs of 0.813 (ER), 0.740 (PR), 0.821 (HER-2), and 0.796 (Ki-67). DCA demonstrated greater net clinical benefit for 3D models, and DeLong’s test revealed statistically significant differences between 2D and 3D models (p < 0.05) for all biomarker groups.

**Discussion:** Both 2D and 3D DCE-MRI radiomics models demonstrated substantial potential for predicting molecular marker expression in breast cancer. However, 3D models exhibited superior performance, likely attributable to their comprehensive capture of spatial tumor heterogeneity. These findings support the preferential use of 3D radiomics models in future research and clinical practice for non-invasive biomarker assessment.

## 1.0 Introduction

### 1.1 Epidemiology and Clinical Significance of Breast Cancer

Breast cancer remains one of the most prevalent malignancies among women worldwide, with continuously rising incidence and diverse clinical manifestations. The American Cancer Society estimated 2.3 million new breast cancer cases globally in 2020, with projections indicating a 0.5% annual increase in both developing and developed countries through 2022. This disease affects postmenopausal women and younger patients alike, significantly impacting quality of life and survival outcomes (1).

### 1.2 Importance of Molecular Biomarkers in Breast Cancer

Estrogen receptor (ER), progesterone receptor (PR), human epidermal growth factor receptor 2 (HER-2), and Ki-67 are molecular biomarkers critical for breast cancer classification and treatment planning. ER and PR expression indicates potential responsiveness to hormone therapy and guides endocrine treatment decisions (2). HER-2 positivity, while associated with more aggressive disease, enables targeted therapy with agents such as trastuzumab, leading to improved outcomes (3). Ki-67, a nuclear proliferation marker, reflects tumor cell proliferation rate; higher levels correlate with higher tumor grade and poorer prognosis (4). Therefore, accurate identification and quantification of these biomarkers are essential in clinical practice for optimizing treatment strategies.

### 1.3 Limitations of Conventional Histopathology

Traditionally, biomarker expression has been assessed using immunohistochemistry (IHC) on tissue samples obtained from core needle biopsy or surgical resection. Although histopathology represents the gold standard, it is invasive, time-consuming, and susceptible to sampling errors due to tumor heterogeneity. Consequently, there is growing interest in imaging-based methods that can predict biomarker expression more reliably and non-invasively.

### 1.4 Role of MRI and Radiomics in Breast Cancer Diagnosis

Magnetic resonance imaging (MRI) has emerged as a highly sensitive imaging modality for breast cancer. Dynamic contrast-enhanced MRI (DCE-MRI) provides superior visualization of tumor margins, assessment of tumor vasculature, and temporal contrast enhancement patterns. DCE-MRI demonstrates a sensitivity of 86–100% in diagnosing breast lesions (5, 6). Radiomics, which converts conventional diagnostic images into quantifiable data, enhances diagnostic precision, treatment modeling, and risk stratification (7, 8).

### 1.5 Current Applications and Knowledge Gaps in Radiomics Studies

Several studies have utilized MRI-based radiomics to predict breast cancer molecular subtypes and biomarkers. For instance, one study demonstrated that integrating DCE-MRI and diffusion-weighted imaging (DWI) radiomics features effectively predicted histologic grade and Ki-67 expression (9). Chiacchiaretta et al. (10) employed standard MRI sequences to predict early recurrence by establishing a radiomic signature. However, many studies have examined only 2D or 3D radiomics without directly comparing the two approaches.

### 1.6 Impact of Dimensionality on Radiomics Performance

The dimensionality of radiomic features whether 2D (extracted from single slices) or 3D (from entire tumor volumes) affects predictive performance. 2D features, extracted from the axial slice with the largest tumor diameter, require less computational time and effort but may omit spatial information critical for accurate modeling. Conversely, 3D features capture the entire tumor volume and may provide more informative and stable results (11, 12). Some studies have reported superior predictive power with 3D models. For example, Wan et al. (13) demonstrated improved discrimination between benign and malignant pulmonary lesions using 3D radiomics. Others argue that the additional computational burden of 3D modelling may not improve performance, particularly for small tumours or when noise and segmentation variability are amplified (14, 15).

### 1.7 Study Rationale and Objectives

Despite these findings, few systematic comparisons of 2D and 3D radiomics models have been conducted to predict a comprehensive panel of breast cancer biomarkers using DCE-MRI. Some studies, such as Zhou et al. (16), used 2D parametric maps to predict HER-2 but excluded ER and PR or did not incorporate 3D features. Additionally, variability in slice thickness, imaging protocols, and manual segmentation affects consistency and comparability across studies (17, 18).

To address these gaps, this study aims to develop and validate 2D and 3D DCE-MRI radiomics models for the non-invasive prediction of ER, PR, HER-2, and Ki-67 expression in breast cancer. This research also seeks to establish a robust methodological framework for future breast cancer radiomics studies and support the clinical implementation of MRI-based biomarker prediction.

## 2.0 Materials and Methods

### 2.1 Study Design and Patient Selection

This post-hoc analysis included 385 female patients with pathologically confirmed breast cancer who underwent DCE-MRI between January 2017 and March 2023, Gathered through written informed consent. Inclusion criteria were: (1) adherence to standardised MRI protocols at participating centres; (2) no prior neoadjuvant chemotherapy at the time of imaging; (3) presence of a solitary mass-type breast lesion on DCE-MRI; and (4) histopathological confirmation of invasive breast cancer with IHC data for ER, PR, HER-2, and Ki-67. Exclusion criteria included incomplete imaging data, multifocal lesions, or absence of molecular marker data. Patient age ranged from 21 to 88 years, with a mean age of 48 years. Patients were stratified into ER+/−, PR+/−, HER-2+/−, and Ki-67 high/low subgroups based on established clinical thresholds. A stratified sampling technique was employed to divide the dataset into training (70%) and validation (30%) cohorts while maintaining biomarker distribution balance. Clinical data and imaging of the 385 patients were retrospectively sourced from Institutional Picture Archiving and Communication Systems (PACS) as well as the electronic medical records on 21/04/2024 for the study purposes.

### 2.2 Immunohistochemistry and Biomarker Classification

Tumor samples were obtained via biopsy or excision and analyzed using IHC according to ASCO/CAP guidelines (2, 19). ER and PR were classified as positive if ≥1% of tumor cells demonstrated nuclear staining. HER-2 status was determined by membrane staining intensity and extent, scored from 0 to 3+. Cases with a 3+ score were considered HER-2 positive; those with 2+ scores underwent fluorescence in situ hybridization (FISH) to confirm amplification, with non-amplified cases classified as HER-2 negative. Ki-67 was classified as high if ≥14% of tumor cells showed positive nuclear staining, based on St. Gallen guidelines (20).

### 2.3 MRI Acquisition Protocol

All patients underwent breast MRI on a GE Signa HDxt 3.0T superconducting magnet system equipped with an 8-channel phased-array breast coil. DCE-MRI employed axial Vibrant+ sequences with the following parameters: repetition time (TR) of 4.3 ms, echo time (TE) of 2.1 ms, slice thickness of 1.6 mm, and slice gap of 0.8 mm. Nine imaging phases were acquired: one pre-contrast phase and eight post-contrast phases. Gadolinium-diethylenetriamine pentaacetic acid (Gd-DTPA) was administered at a dose of 0.1 mmol/kg body weight via power injector at a flow rate of 2.0 mL/s, followed by a 20 mL saline flush. Post-contrast images were acquired immediately and at intervals from 61 to 429 seconds following contrast administration.

### 2.4 ROI Segmentation and Image Preprocessing

Image preprocessing and lesion segmentation were performed using the Reforma program on the GE AW 4.6 workstation. Axial images from the third post-contrast phase were selected due to optimal tumour enhancement. Slice thickness and spacing were resampled to 0.8 mm using the Loop program. Lesions were delineated using a semi-automated method in the Auto Contour tool, which employed a dynamic threshold region-growing algorithm. An operator positioned the cursor at the lesion centre, and the software automatically generated a 3D volume of interest (3D-ROI). For 2D segmentation, the tumour was outlined on the axial slice with the largest tumour diameter, designated as the 2D region of interest (2D-ROI). Contours were refined using the “Cut Outside” tool to exclude non-tumour tissue. When automatic segmentation was suboptimal, the process was repeated with adjusted parameters. Final ROIs were exported in NIfTI (.nii) format using ITK-SNAP software version 4.02 for radiomic analysis.

### 2.5 Radiomic Feature Extraction and Selection

Radiomic feature extraction from both 2D-ROIs and 3D-ROIs was performed using Intelligence Foundry (IF) 3.1 software (GE Healthcare). Extracted features included first-order statistics, second-order histogram-based features, and higher-order textural features derived from grey-level co-occurrence matrix (GLCM), grey-level run length matrix (GLRLM), and wavelet transforms. A total of 1,427 radiomic features were extracted per lesion.

Several preprocessing steps were implemented to reduce dimensionality and enhance generalization. First, features with near-zero variance were removed. For normally distributed features, Pearson correlation coefficients were calculated; for non-parametric features, Spearman correlation was used to eliminate highly collinear variables (r > 0.9). To mitigate the influence of outliers, the top and bottom 5% of values in both training and validation datasets were replaced using the formula: [0.9 × Percentile + 0.1 × random() × (Percentile[1] − Percentile[0])]. All features were then standardized using z-score normalization based on training set means and standard deviations.

LASSO regression with 10-fold cross-validation was subsequently applied to identify the most predictive features. Multivariable logistic regression models were constructed for both 2D and 3D datasets using the selected features for each biomarker.

### 2.6 Model Construction and Evaluation

Logistic regression models were developed independently for ER, PR, HER-2, and Ki-67 using 2D and 3D radiomic features. All models were trained on the 70% training cohort and tested on the 30% validation cohort. Model performance was assessed using receiver operating characteristic (ROC) curve analysis. Performance metrics included area under the curve (AUC), sensitivity, specificity, accuracy, positive predictive value (PPV), and negative predictive value (NPV). Decision curve analysis (DCA) was performed to evaluate the net clinical benefit of each model across different threshold probabilities. DeLong’s test was employed to statistically compare the AUCs of 2D and 3D models for each biomarker.

### 2.7 Statistical Analysis

All statistical analyses were conducted using SPSS Statistics 26.0 (IBM Corp., Armonk, NY, USA) and R version 3.6.1. The Shapiro-Wilk test was used to assess the normality of continuous variables. Normally distributed continuous data were compared using independent sample t-tests, while non-normally distributed data were analyzed using the Mann-Whitney U test. DeLong’s test was applied to compare the AUCs of paired 2D and 3D models. A two-tailed p-value < 0.05 was considered statistically significant.

Sample size was calculated to detect a significant difference in AUC between 2D and 3D radiomics models using DeLong’s test (α = 0.05, power = 80%). Assuming a moderate AUC difference of 0.08, a correlation of 0.8 between 2D and 3D model predictions due to shared imaging data, and a 70%/30% training/validation split, a minimum of 300 patients (90 in validation) was required. The study enrolled 385 patients (116 in validation), exceeding this requirement and ensuring adequate statistical power. Assumptions included normality of AUC differences, independence of patient data, and balanced biomarker subgroups achieved through stratified sampling.

## 3.0 Results

### 3.1 ROI Segmentation and Radiomics Workflow

The workflow for patient-specific ROI delineation is depicted in **Figure 1**, illustrating the generation of 2D and 3D ROIs using a semi-automated segmentation technique. The 2D-ROI was derived from the slice demonstrating the maximum tumor diameter, while the 3D-ROI was reconstructed from the entire lesion volume, providing comprehensive spatial information regarding contrast enhancement distribution. Figure 2 illustrates the complete radiomics workflow, encompassing ROI delineation, feature extraction, feature selection, and model construction.

**Figure 1:**
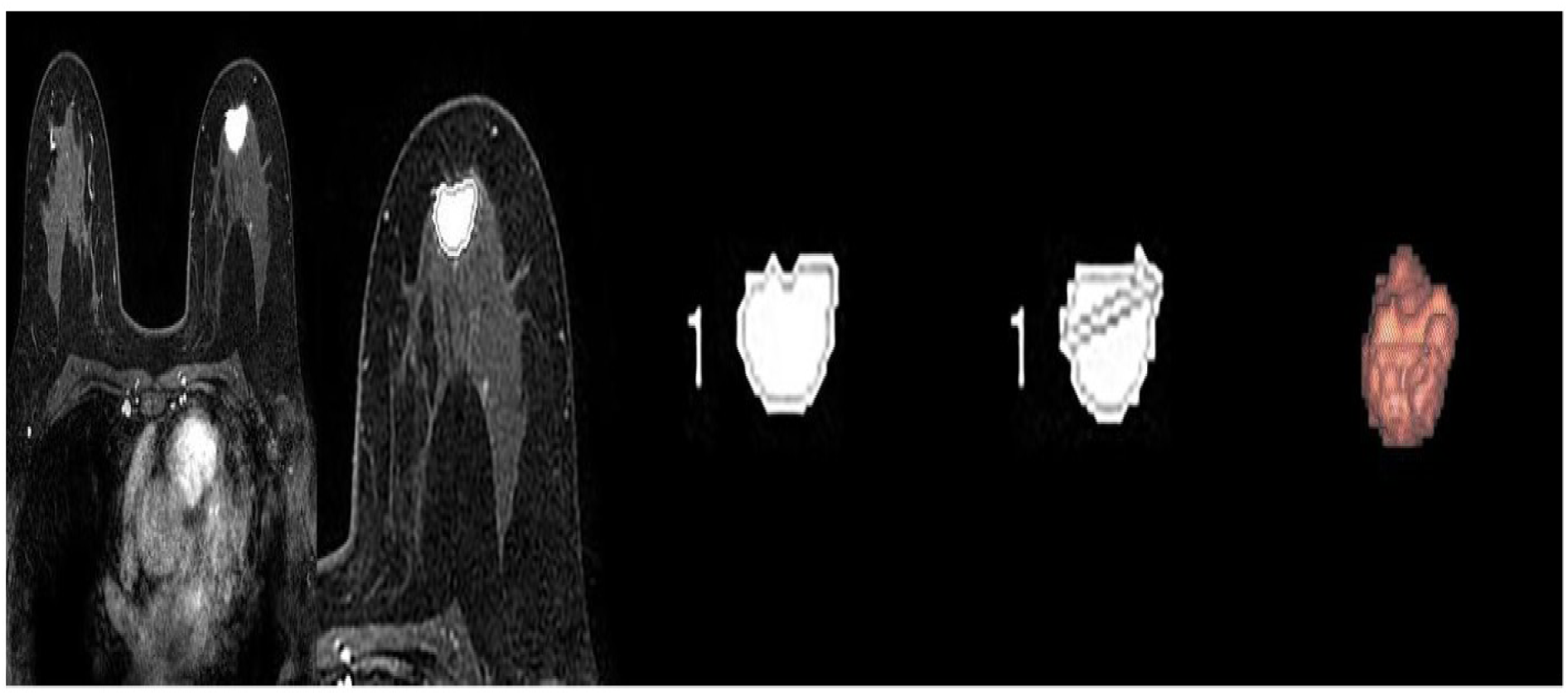
ROI delineation for breast cancer. (a) Phase 3 DCE-MRI image showing marked enhancement in the upper inner quadrant of the left breast. (b) Automatic ROI delineation following manual selection of the most enhanced area (white circle). (c) Segmented ROI after background removal. (d) The slice with maximum tumor diameter identified automatically, marked by a diagonal line representing the 2D-ROI. (e) Red pseudo-colored image visualizing the 3D-ROI.

**Figure 2:**
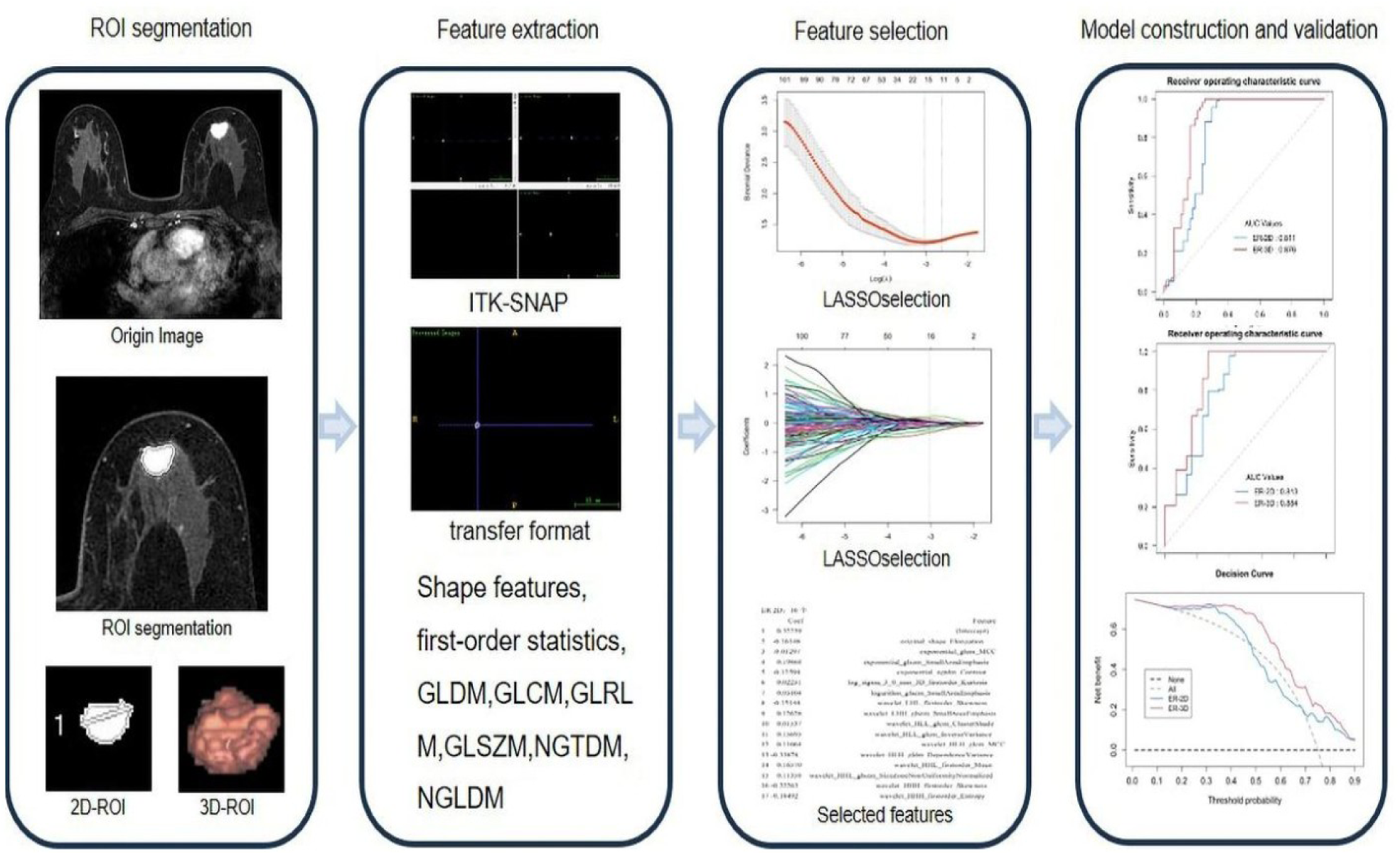
Radiomics analysis workflow. The process includes delineation of 2D-ROI and 3D-ROI via interactive semi-automatic segmentation, importing ROIs into ITK-SNAP for format conversion, extracting high-throughput features from both ROI types, performing feature selection, and constructing and validating predictive models.

### 3.2 Radiomic Feature Selection Using LASSO

Both 2D and 3D radiomic features were subjected to LASSO regression with cross-validation for optimal feature selection. Figure 3 depicts the LASSO analysis for each biomarker. The final selected features were: 16 for ER-2D and 18 for ER-3D; 10 for PR-2D and 8 for PR-3D; or HER-2-2D and 10 for HER-2-3D; and 13 for Ki-67-2D and 11 for Ki-67-3D. Notably, 3D models retained a greater number of relevant features in several cases.

**Figure 3:**
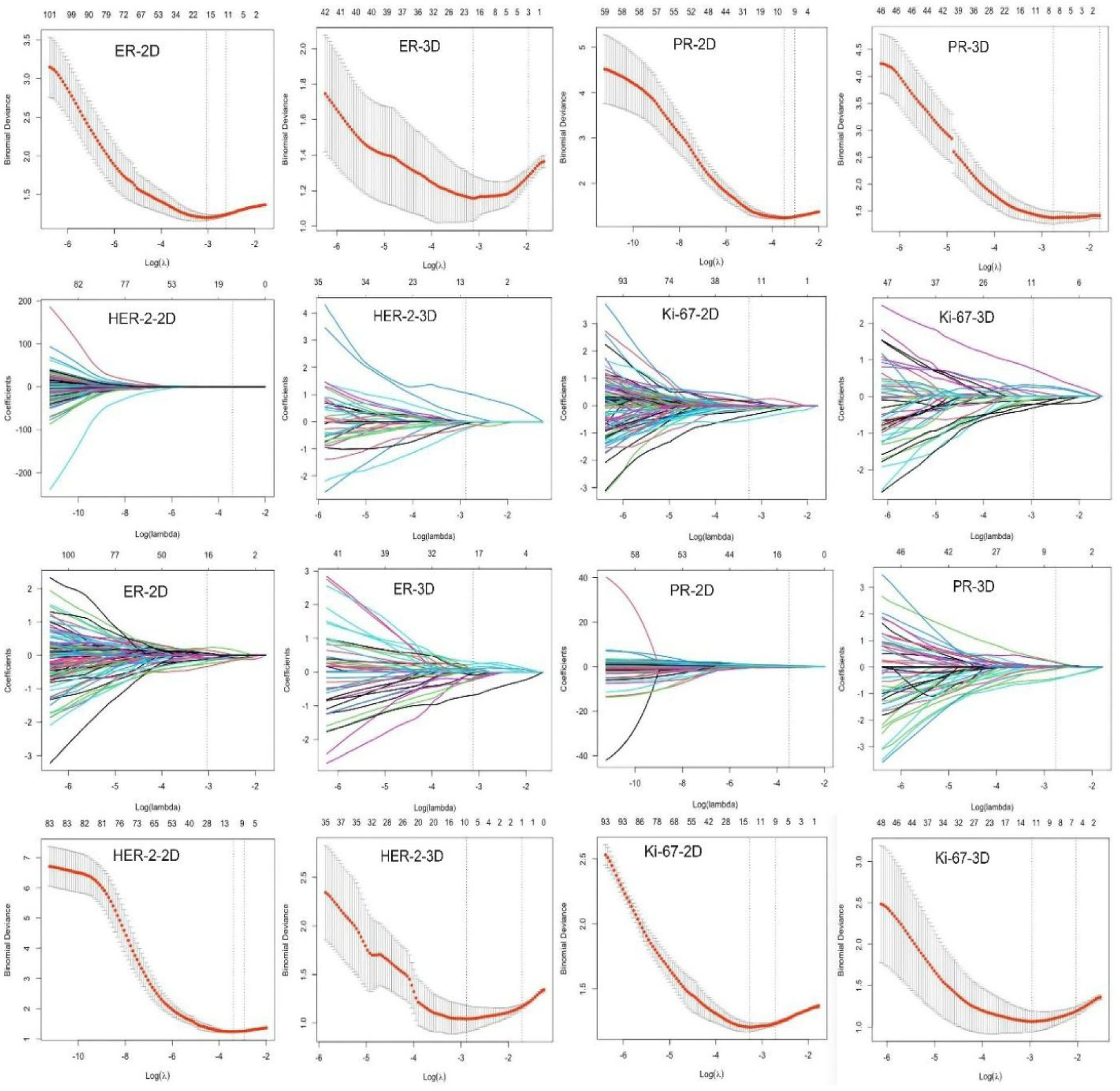
LASSO algorithm for feature selection. (a1-8) Through 10-fold cross-validation, the parameter λ is optimized to minimize the binomial deviance, removing redundant features. (b1-8) Coefficients are converged using one-way analysis of variance to select optimal features. Final selected features: ER-2D (16), ER-3D (18), PR-2D (10), PR-3D (8), HER-2-2D (12), HER-2-3D (10), Ki-67-2D (13), and Ki-67-3D (11).

### 3.3 Model Performance Metrics

Table 1 summarizes the performance metrics for all biomarker predictive models. Across all biomarkers ER, PR, HER-2, and Ki-67 the 3D models consistently demonstrated superior performance compared to 2D models. For ER, the 3D model achieved a validation AUC of 0.884, outperforming the 2D model (AUC = 0.813). Similarly, for PR, the 3D model attained an AUC of 0.861 versus 0.740 for the 2D model. HER-2 models showed comparable trends, with AUCs of 0.898 (3D) and 0.821 (2D). For Ki-67, the 3D model achieved an AUC of 0.886 compared to 0.796 for the 2D model.

For HER-2 prediction, the 2D model achieved training and validation accuracies of 0.830 and 0.821, respectively, while the 3D model attained accuracies of 0.907 and 0.898. The superior accuracy of the 3D model for HER-2 is consistent with previous research indicating that HER-2-positive tumors exhibit greater spatial heterogeneity, which is better captured by volumetric analysis. Similar trends were observed for Ki-67 models. The 2D model achieved training and validation AUCs of 0.749 and 0.796, respectively, whereas the 3D model achieved higher AUCs of 0.874 and 0.886. The 3D model for Ki-67 also demonstrated improved specificity and positive predictive value, suggesting enhanced capability to identify high-proliferation tumors.

### 3.4 Statistical Comparison Using DeLong’s Test

DeLong’s test was employed to statistically compare the AUCs of 2D and 3D models. Table 2 demonstrates that p-values for all four biomarkers were statistically significant in both training and validation sets, with p-values ranging from 0.00135 to 0.03082 in the training set and 0.00322 to 0.02657 in the validation set. These results confirm that 3D radiomic models provide superior representation of tumor biology compared to 2D models.

**Table 2:**
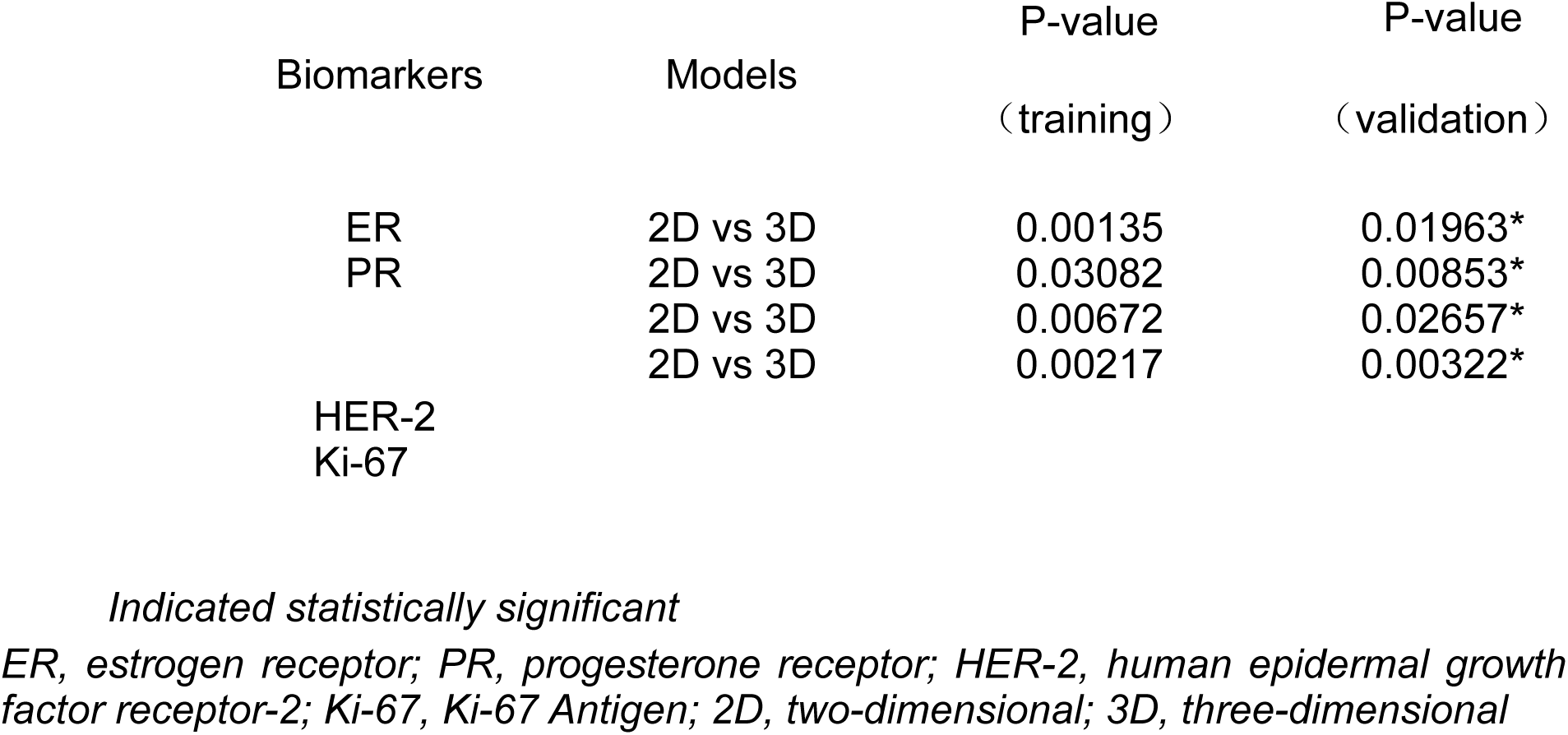
DeLong’s Test Results for AUC Comparison.

### 3.5 ROC Curve Analysis

Figure 4 presents ROC curves for each biomarker in both training and validation sets. The curves visually demonstrate larger AUCs for 3D models, corroborating the quantitative data in Table 1. Figure 4 displays the ER, PR, HER-2, and Ki-67 ROC curves for both 2D and 3D models in training (a1-4) and validation (b1-4) sets. Both model types demonstrated strong performance, with consistently higher AUC values for 3D models compared to their 2D counterparts.

**Figure 4:**
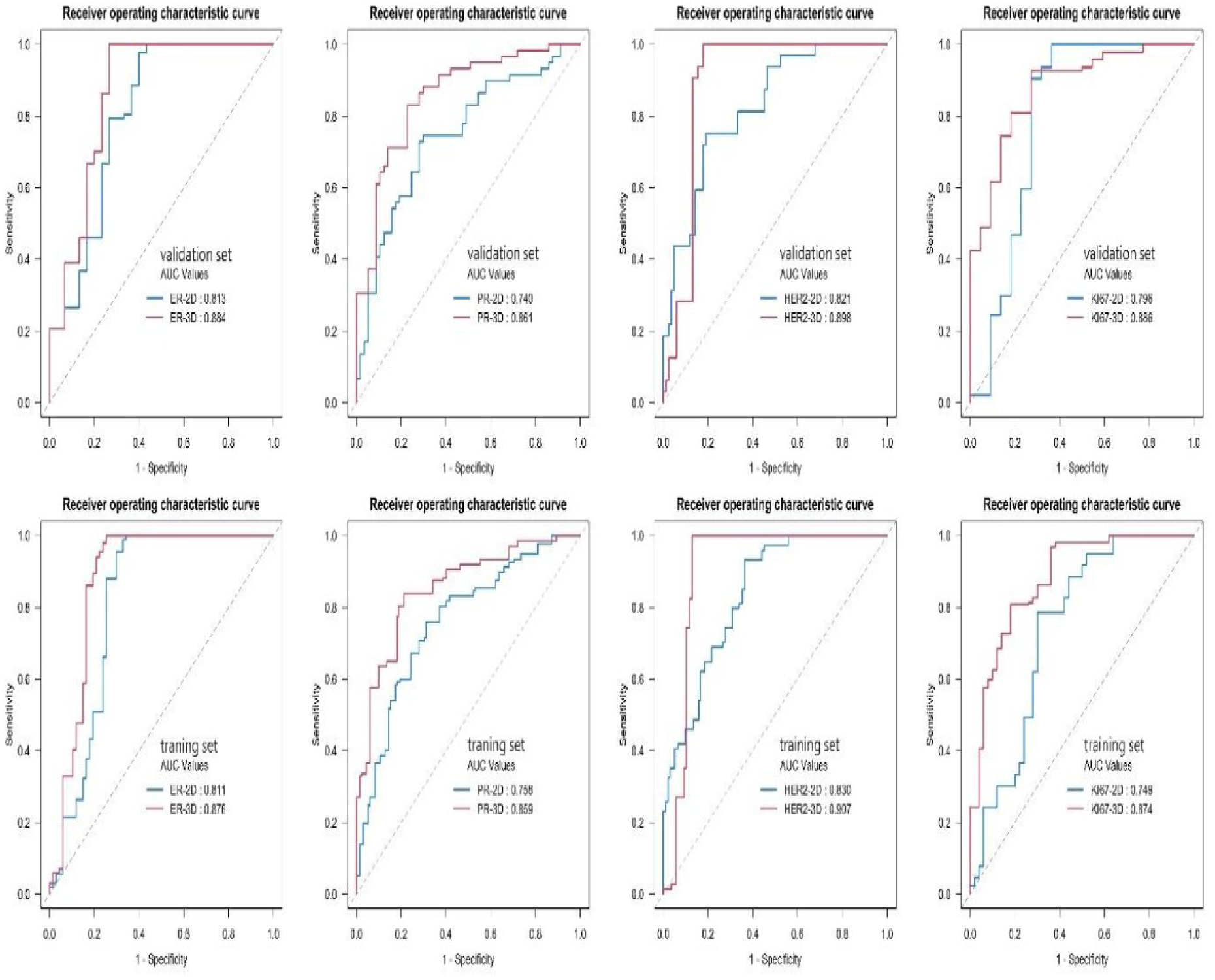
ROC curves for ER, PR, HER-2, and Ki-67 prediction using 2D and 3D models. (a1-4) Training set. (b1-4) Validation set. Both models demonstrated excellent performance, with 3D models consistently achieving higher AUC values than corresponding 2D models.

### 3.6 Clinical Utility Assessed via Decision Curve Analysis

Decision curve analysis (DCA) was performed to evaluate the net clinical benefit of each model across various threshold probabilities. Figure 5 illustrates that the 3D models consistently provided higher net benefit at nearly all threshold levels compared to 2D models. This reinforces the hypothesis that 3D models are more clinically useful for decision-making. For example, HER-2 prediction demonstrated significantly higher net benefit in the 10–70% threshold probability range, which is particularly relevant for therapeutic decision-making.

**Figure 5:**
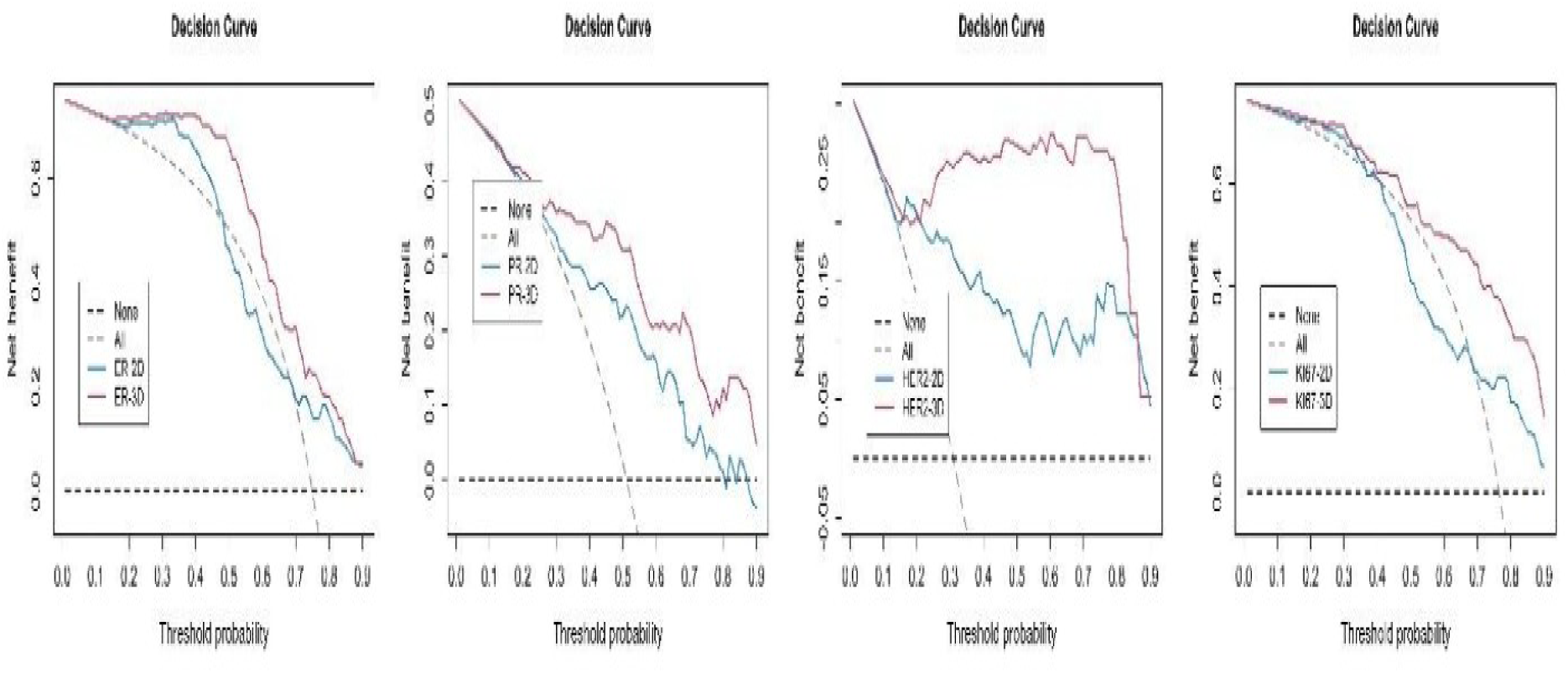
Decision curve analysis comparing 2D and 3D models for ER (a1), PR (a2), HER-2 (a3), and Ki-67 (a4). The x-axis represents threshold probability and the y-axis represents net benefit. The findings revealed that 3D models in each group demonstrated superior net benefit compared to their respective 2D counterparts.

### 3.7 Interpretation and Clinical Implications

The consistently superior performance of 3D models across all biomarkers indicates that volumetric feature representation conveys more comprehensive spatial information related to tumor heterogeneity. The enhanced performance of 3D models can be attributed to their capacity to incorporate axial (z-axis) data regarding slice-to-slice variations in tumor morphology and contrast enhancement kinetics characteristics that likely reflect underlying molecular features. Furthermore, the semi-automated segmentation approach minimizes inter-observer variability and facilitates efficient generation of 3D-ROIs within clinically feasible timeframes.

These findings have significant clinical implications. Accurate non-invasive biomarker prediction based on 3D radiomics data may reduce the need for repeated biopsies and optimize surgical planning. The improvements in HER-2 and Ki-67 predictive models demonstrate potential applications in selecting neoadjuvant treatment regimens or identifying candidates for clinical trials.

## 4.0 Discussion

This study systematically compared the performance of 2D and 3D radiomics models derived from DCE-MRI for predicting breast cancer molecular biomarkers, including ER, PR, HER-2, and Ki-67. The results consistently demonstrated that 3D models exhibited superior discriminatory power, clinical net benefit, and statistical significance compared to 2D models. These findings suggest that volumetric radiomics analysis (VRA) may provide a more reliable and clinically valuable non-invasive alternative to immunohistochemical assessment for molecular characterization of breast cancer.

The superiority of 3D models supports the theoretical premise that volumetric features more effectively capture tumor heterogeneity at both genetic and phenotypic levels. Tumor heterogeneity has been shown to contribute to therapy resistance and poor prognosis in breast cancer patients (21). Radiomic features derived from entire tumor volumes better reflect variations in pixel intensity, enhancement patterns, and morphological complexity that cannot be captured by a single axial slice at the tumor’s maximum diameter. Previous studies examining spatial heterogeneity through imaging have demonstrated that complexity measures such as entropy and fractal dimension correlate with histological invasiveness and molecular biomarkers (22, 23). Therefore, the improved accuracy of 3D models may stem from their enhanced ability to capture biologically relevant imaging signatures.

This study employed a semi-automatic segmentation method using dynamic threshold region-growing to minimize bias and improve ROI delineation accuracy for both 2D and 3D imaging. ROI segmentation accuracy directly influences feature extraction and subsequent model performance. Manual contouring methods are often time-consuming and subject to inter-observer variability, particularly when tumor margins are poorly defined. Automated or semi-automated segmentation has been shown to enhance feature reproducibility and reduce operator-induced variability (24, 25). Additionally, applying consistent 3D-ROIs using the proposed semi-automated technique helps overcome a major barrier to clinical translation of 3D radiomics: the time-intensive nature of volumetric annotation.

A methodological strength of this study is the use of a standardized imaging protocol with uniform slice thickness of 0.8 mm. Previous research has demonstrated that variations in imaging parameters such as slice thickness, reconstruction algorithms, and contrast timing affect radiomic feature reproducibility (26). Variability of features across institutions and scanners represents a major challenge in radiomics, complicating clinical implementation. By employing a homogeneous imaging protocol, this study minimized methodological variance and strengthened internal validity for comparing 2D and 3D models.

Biomarker-specific analysis revealed particularly significant improvements in predictive accuracy for 3D models of HER-2 and Ki-67. HER-2 overexpression is associated with increased angiogenesis and perfusion, resulting in elevated DCE-MRI signal intensity and characteristic kinetic patterns (27). Ki-67, as a proliferation marker, correlates with invasive tumor margins and internal heterogeneity parameters that can be identified through volumetric texture analysis (28). While previous studies using perfusion-weighted and diffusion-weighted imaging have linked imaging-derived heterogeneity to Ki-67 status, none have systematically compared 2D and 3D radiomics across a comprehensive panel of biomarkers as performed in this study.

Despite strong evidence supporting the clinical utility of 3D radiomics, this approach introduces higher feature dimensionality. High-dimensional feature spaces can lead to model overfitting, particularly with small sample sizes (29). To mitigate this risk, LASSO regression combined with cross-validation was employed for feature selection—recognized as an effective strategy for enhancing model performance in high-dimensional contexts (30). Nevertheless, external validation using independent datasets is essential for evaluating the generalizability of these models in diverse clinical settings.

An expanding body of literature supports the integration of radiomics with genomics, proteomics, and clinical variables to enhance predictive performance—a concept termed radiogenomics, which is increasingly applied in oncology (31). Combining 3D radiomics with molecular assays such as Oncotype DX or MammaPrint could further improve biomarker-based risk stratification and reduce dependence on invasive biopsies, particularly for multifocal or anatomically challenging lesions. Future research should adopt such integrative frameworks using multimodal machine learning (MML) pipelines.

Beyond scientific considerations, computational resource requirements merit discussion. While 3D modeling is more computationally intensive than 2D modeling in terms of processing power and storage, advances in cloud computing and GPU-accelerated pipelines have made such analyses increasingly feasible (32). Moreover, time savings achieved through semi-automatic ROI delineation may offset the increased processing time for 3D analysis, making it a potentially practical option for routine clinical use.

This study has several limitations. First, it was retrospective, cross-sectional, and conducted at a single center, which may limit generalizability. Although the results represent reproducible, clinically valid models, they were not derived from heterogeneous, multi-institutional datasets. Second, while DCE-MRI provided excellent soft tissue contrast and functional information, the absence of diffusion-weighted or T2-weighted sequences may have limited diagnostic comprehensiveness. Multimodal radiomics incorporating multiple MRI sequences could further enhance model performance. Finally, no biological validation was performed, such as correlating radiomic features with histological architecture; future research should pursue prospective validation with histopathological correlation.

In conclusion, 3D radiomics models based on DCE-MRI significantly improve biomarker prediction accuracy in breast cancer compared to 2D models. With appropriate data management and robust feature selection, 3D radiomics has the potential to become a cornerstone of precision imaging biomarkers, reducing reliance on invasive biopsies and supporting personalized treatment planning. Continued research is essential as radiomics transitions from a research tool to a clinically validated technology, particularly when integrated with biological and clinical data.

## 5.0 Conclusion

Both 2D and 3D DCE-MRI-based radiomics models demonstrate excellent performance in accurately predicting the expression status of molecular markers ER, PR, HER-2, and Ki-67 in breast cancer. The superior performance of 3D models over 2D models supports their preferential use in future breast cancer radiomics research. However, it is essential to employ appropriate methodologies to address the additional workload associated with 3D-ROI delineation and minimize noise interference. These findings underscore the significant clinical potential of 3D radiomics models in preoperative treatment planning and personalized therapy for breast cancer patients.

## 6.0 Declarations

### Ethical Approval and Consent to Participate

This study was approved by the First Affiliated Panzhihua Municipal Central Hospital Medical Ethics Committee (Approval/Reference No. **PKLS Zi NO. [2024-005]**) and was conducted in accordance with the principles of the Declaration of Helsinki. Written informed consent was obtained from all participants before enrollment, and all patient data were anonymized to ensure confidentiality.

### Consent for Publication

All authors have consented to submit and publish this manuscript in BMC Medical Imaging.

### Availability of Data and Materials

All data and materials are included in the manuscript.

### Authors’ Contributions

All authors declare that they have made equal contributions to this work.

## Abbreviations

2D: Two-dimensional
3D: Three-dimensional
ASCO/CAP: American Society of Clinical Oncology/College of American Pathologists
AUC: Area under the ROC Curve
BC: Breast Cancer
CI: Confidence Interval
DCA: Decision Curve Analysis
DCE-MRI: Dynamic Contrast-Enhanced Magnetic Resonance Imaging
DWI: Diffusion-Weighted Imaging
ER: Estrogen Receptor
FISH: Fluorescence In Situ Hybridization
Gd-DTPA: Gadolinium-Diethylenetriamine Pentaacetic Acid
GLCM: Gray Level Co-occurrence Matrix
GLRLM: Gray Level Run Length Matrix
HER-2: Human Epidermal Growth Factor Receptor 2
IF: Intelligence Foundry
IHC: Immunohistochemistry
Ki-67: Ki-67 Proliferation Marker
LASSO: Least Absolute Shrinkage and Selection Operator
MML: Multimodal Machine Learning
MRI: Magnetic Resonance Imaging
NIfTI: Neuroimaging Informatics Technology Initiative
NPV: Negative Predictive Value
PPV: Positive Predictive Value
PR: Progesterone Receptor
ROC: Receiver Operating Characteristic
ROI: Region of Interest
TE: Echo Time
TR: Repetition Time
VRA: Volumetric Radiomics Analysis

## Acknowledgements

Not applicable.

## Supporting information

Figure 1: The delineation of the ROI for breast cancer. a Exhibits the phase 3 DCE image, revealing significant enhancement in the upper inner quadrant of the left breast. Demonstrates automatic ROI delineation after manually selecting and marking the area with the most pronounced enhancement using a white circle. c Displays the segmented ROI following background removal. d represents the layer where the tumor’s maximum diameter is automatically identified, indicated by a diagonal line as a two-dimensional 2D-ROI. Presents a red pseudo-colored image to visualize the three-dimensional 3D-ROI.

Figure 2: The workflow of radiomics analysis. Outlining 2D-ROI and 3D-ROI by interactive semi-automatic segmentation. The obtained ROIs were imported into ITK-SNAP to convert the transfer format and extract high-throughput features based on two different ROIs. Feature selection. Model construction and validation

Figure 3: The LASSO algorithm filters imaging omics features to select optimal features.

a(1–8) Through 10-fold cross-validation, parameter λ is adjusted, and the model’s fitting loss binomial bias is minimized, resulting in the removal of redundant imaging omics features.

b(1–8) One-way analysis of variance is then employed to converge coefficients and select the optimal imaging omics features. The final selected features included ER-2D 16, ER-3D 18, PR-2D 10, PR-3D 8, HER-2-2D 12, HER-2-3D 10, Ki-67-2D 13, and Ki-67-3D 11

**Table 1**: Comparative Performance of 2D and 3D Models ER, estrogen receptor; PR, progesterone receptor; HER-2, human epidermal growth factor receptor-2; Ki-67, Ki-67 Antigen; 2D, two-dimensional; 3D, three-dimensional; AUC, area under the ROC curve; CI, confidence interval; PPV, positive predictive value; NPV, negative predictive value

**Table 2**: DeLong’s Test Results for AUC Comparison *ER, estrogen receptor; PR, progesterone receptor; HER-2, human epidermal growth factor receptor-2; Ki-67, Ki-67 Antigen; 2D, two-dimensional; 3D, three-dimensional*

Figure 4 presents the ER, PR, HER-2, and Ki-67 ROC curves for both 2D and 3D models. A (1–4) The training set. b (1–4) The validation set. Both models demonstrated exceptional performance, with higher AUC values for the 3D models than their corresponding 2D models.

Figure 5. The DCA of 2D and 3D models for ER(a1), PR(a2), HER-2(a3), and Ki-67(a4) was compared, with the x-axis representing threshold probability and the y-axis representing net profit. The findings revealed that the 3D models in each group demonstrated *superior profitability compared to their respective 2D counterparts*

